# Motor Equivalence in Motor Awareness

**DOI:** 10.1101/2024.10.29.620842

**Authors:** J Fasola, S Betka, N Faivre, O Blanke, OA Kannape

**Author notes:** contributed equally (O Blanke, OA Kannape).

## Abstract

Motor awareness (MA) describes the level of conscious access we have to the details of our movements and as such is critical to the feeling of control we maintain over our actions (sense of agency). Although our movements rely on specific sensorimotor transformations as well as distinct reference frames for the active body part, or effector, numerous studies report spatiotemporal thresholds of MA that are comparable across different effectors and tasks as well as supramodal. However, this has not been tested directly and there is currently no direct empirical support for effector-independent MA nor a description of a potentially shared underlying mechanisms. We therefore designed a goal-directed reaching paradigm that participants performed once with their upper-limbs, using a joystick, and once with their full body, by leaning and thereby displacing their center-of-mass. We assessed both MA and corrective movements for sensorimotor mismatches by providing either veridical feedback or introducing random spatial deviations. We hypothesized that changes in motor compensation and awareness, across effectors, with and without a concurrent cognitive load, would follow the same pattern of behavior if they relied on a shared underlying mechanism. Our results lend support for such an effector-independent mechanism, as we observed that MA was comparable across effectors: i) in un-deviated control trials with and without cognitive load, ii) in converging trials where the direction of the deviation corresponded to the direction of the target location, and iii) based on strongly correlated psychometric MA thresholds across effectors. At the same time, data from diverging trials, where the direction of the deviation opposed the direction of the target location, indicate that in case of conflicting information and increased kinematic task demands MA draws on effector-specific sensorimotor information, corresponding to performance differences between hand and full body movements observed in baseline blind-reaching versus visually guided reaching. Overall, our findings provide a direct link between low-level sensorimotor transformations and abstract motor representations and their role in MA, consolidating a gap in conceptual frameworks of the sense of agency.

## 1 Introduction

We perform most of our daily activities without paying particular attention to the details of how an action is performed. When we prepare a cup of tea, perhaps the only aspect we consciously monitor is when we pour the hot water into the cup. However, we are not generally aware of the particular grasp we choose to open the cupboard, nor of postural adjustments such as when we shift our weight onto one leg in order to reach for the cup. Nonetheless, we maintain a sense of agency for (SoA) and control over all of the aforementioned actions (even if this sensation is “phenomenally thin” ^1,2^). It has been argued that motor awareness (MA), in other words our ability to consciously monitor the details of our actions, provides one potential limit of our SoA ^3,4^; our SoA should not be more sensitive to perturbations than our ability to perceive and automatically adjust for them. Furthermore, a central underlying mechanism, from finger-related movements to full-body actions, has been proposed, making MA effector-independent ^4–6^. This is in-line with recent evidence for an abstract action system that suggests effector-independent representations for e.g., reaching or grasping movements ^7^, extending the general notion of effector-independent motor representation ^8,9^, to effector-specific sensorimotor transformations. Here, we explicitly tested the effector-independence hypothesis for MA, and by extension SoA, by asking participants to the same, goal-directed reaching task with (1) their right hand (effector task 1) and (2) by leaning with their trunk (effector task 2). We hypothesized that MA thresholds would co-vary between body-part and full-body actions, despite their vastly different skeletomotor organization and different cortical and subcortical control mechanisms ^10,12,14^. To evaluate the robustness of effector-independent MA, we carried out two further experiments. First, we controlled for general differences in reaching performance between the hand and the trunk by asking participants to perform a blind reaching paradigm with the respective effector. Second, we introduced a dual task paradigm, to evaluate MA under cognitive loading; changes induced in MA and motor control, should be reflected and comparable across effectors, if the effector-independent hypothesis is correct.

Action monitoring and MA have been studied since the early work by Nielsen (1963, 1978). These studies have been primarily focused on reaching movements performed with the arms ^3,11,13,15^ and hands ^16,17^, or fingers_^18^. MA is generally evaluated in response to sensorimotor mismatches between visual and proprioceptive feedback and the motor commands and studies consistently demonstrated that participants attributed feedback that was deviated in space by up to 6.5°-15° to their own movement ^15,19–21^, and even larger deviations in neurological patients ^22^ and individuals diagnosed with schizophrenia or with a genetic predisposition for psychosis ^16,23–26^. Similarly, experiments with temporal delays between the participant’s actual hand and the visual feedback illustrated that participants self-attributed movements with up to 150-200ms of delay ^23,27^.

Aside from brief reaching movements, MA and the SoA have further been investigated for continuous movements such as finger-tapping using auditory feedback ^18,28^, or hand-opening and closing using visual feedback ^29,30^. Moreover, footsteps generated during walking, evoke similar psychometric responses in MA as reported for auditory feedback for upper-limb movement ^6^. More recently the research of Kannape et al. ^4,31^ extended these paradigms by studying walking agency using visual feedback. The thresholds in these studies are again comparable to those involving only upper-limb movements both with respect to temporal and spatial characteristics. Taken together, these studies suggest that MA is independent of the end-effector as well as supramodal ^32^, arguing for a common action monitoring mechanism ^6,33^.

MA is intrinsically linked to one’s actual, ongoing movement and its accompanying reafferences ^34^ rather than a mediated action consequence such as an audio-visual stimulus generated via a button press, as commonly used in SoA paradigms. MA has therefore closely been associated with computational models of sensorimotor control, most notably the comparator framework ^35^. The framework relies on a comparison of reafferent feedback and a predicted future state to improve sensorimotor control. It further proposes that small movement errors are automatically corrected via this comparison, as has for example been observed when participants update their movement towards a target object displaced during a saccade ^36^. Larger errors on the other hand may be perceived by participants and consciously corrected as seen when spatiotemporal deviations exceed the MA threshold ^3,11,31^. Further evidence for the comparator and more generally the predictive coding framework ^37^, comes from research on sensory attenuation ^38^, which, relevant to the current study, has similarly been shown to be comparable for action consequences generated by both the hand and the foot ^39^. Therefore, this framework provides a model for online movement adjustments, both automatic and conscious, and sensory attenuation, but also a model mechanism to discriminate between our own actions from the ones generated by the environment ^40^ or other agents ^20^. As this framework applies to most types of movements it also, in principle, supports the idea of effector-independent MA. However, no prior research has investigated MA for comparable movements achieved with two different effectors within a single study and cohort to test this hypothesis.

While co-varying MA thresholds across effectors are one indication that MA may rely on abstracted, effector-independent motor representations, we set out to “stress-test” this hypothesis by running the same paradigm while taxing cognitive resources. Cognitive loading, induced by asking participants to perform a secondary motor or cognitive task in addition to a primary one, has systematic effects on motor control ^41^. Dual-tasking is regularly applied in clinical populations ^42^ and the elderly ^43^ where particularly strong influences have been demonstrated on balance and locomotion. Previous research has also indicated an increased separation between MA and motor performance during dual-tasking ^4,5,44^. During goal-directed walking this was reflected in the finding that cognitive loading did not alter the locomotor trajectories, but impaired motor control (walking velocity) in general, but MA only in trials with high perceptual uncertainty. Understanding how MA is altered during dual-tasking is not only relevant for clinical application but also may shed further light on how it differs for various types of movements and effectors. For that reason, a visual color-word Stroop task (Stroop 1935) was introduced for the movement of the full-body and the hand in this study.

Finally, potential effector-dependent differences in MA relating to the spatial mismatches investigated here could also be due to baseline differences in reaching performance. E.g., higher MA thresholds for one effector may reflect an increased difficulty to perform a motor task, rather than a difference in the MA mechanism. In order to control for differences in hand- and trunk-based reaching, we asked participants to perform blind reaching tasks both with their upper-limb and by leaning with their full-body. This is further motivated by a recent study suggesting differences in error and variability between the two effectors ^45^, suggesting that trunk-controlled movements may be better than hand-controlled movements, even though the latter are more common with respect to visually guided feedback.

The primary goal of this study was thus to test the hypothesis that MA is effector-independent and therefore comparable between goal-directed movements of the hand (reaching) and the full-body (leaning). We hypothesized that MA thresholds would co-vary between the two effectors suggesting that MA may be based on an abstracted motor representation as opposed to the effector-specific sensorimotor transformations. In addition to the visually guided movements of the MA task, we further controlled for baseline differences in sensorimotor control between the hand and trunk in a blind reaching task. Finally, we investigated the effects of cognitive loading on MA and movement kinematics. If MA is modulated by cognitive loading, we hypothesized to observe comparable changes for both effectors in relation to changes in motor performance.

## 2 Results

### 2.1 Action monitoring

Motor awareness (MA) was defined as the ratio of yes-responses out of all trials of the same deviation. In a first step, we evaluated if MA was affected by independent variables linked to the setup, namely the location of the targets or the direction of the deviation. While no significant effect was observed for Target Side (p=0.07), there was a strong interaction between Target Side and Deviation Side (i.e., sign of deviation, p<0.001). MA was less sensitive, that is, yes-responses in deviated trials were significantly more frequent, when the deviation side converged towards the target side. In other words, fewer experimentally induced deviations were noticed, when motor compensation was towards the center or midline. Given this interaction, we did not collapse the trials across targets and absolute deviation (cf. ^5,31,44^). Instead, as the effect was symmetrical around both targets, we grouped the trials by mode: *diverging*, for negative or conflicting deviations, and *converging*, for positive or “helpful” deviations; control trials with no deviation were analyzed separately.

MA ratings in control trials with no deviation should be high, as participants receive veridical feedback of their movements ^33^. Based on previous studies this correct MA should also be robust to cognitive loading ^5,44^. In-line with these hypotheses, participants correctly self-attributed 89.9 ± 1% and 91.3 ± 1% (mean ± sem) of the non-deviated trials for the full body and hand, respectively (Figure 2D, shaded area). There is strong evidence in favour of H_0_, i.e., that these distributions do not differ between the two effectors (BF_10_ = 0.07, paired T-test: p = 0.25). We neither observed an effect of Task or Effector, nor an interaction (p>0.43; see Figure 3E). As before, we used Bayesian t-tests to show strong evidence in favour of H_0_ (overall BF_10_<0.18, paired T-test: p>0.11): MA in control trials does not differ across effectors.

**Figure 1.**
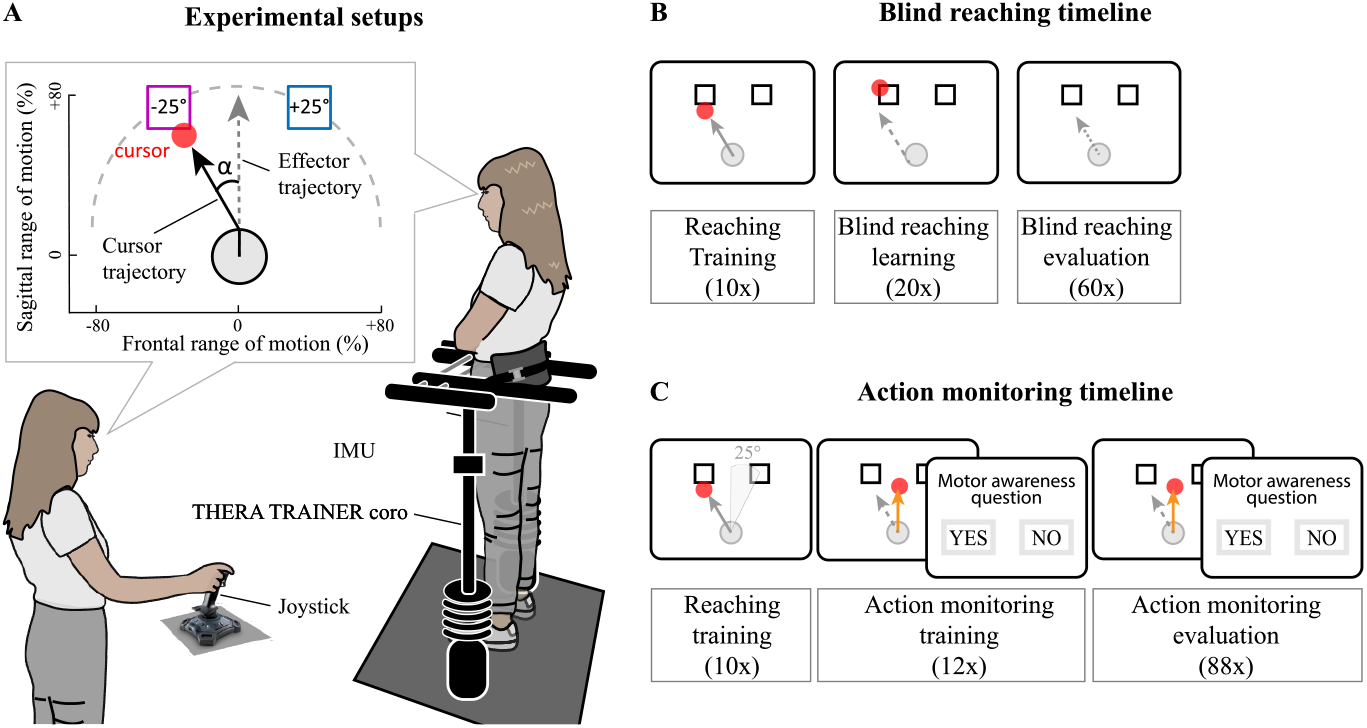
**A**. Experimental setup and visual feedback. Participants either controlled the cursor by leaning their whole body inside the Thera Trainer Coro or by moving the joystick with their hand (counter-balanced). In the action monitoring task, the cursor trajectory could be deviated by an angle (α) that participants had to compensate in order to reach the target. **B**. Baseline, blind reaching task. Participants started the experiment with 10 training trials with continuous, congruent visual feedback. They were then trained to reach the target with only terminal feedback. During the evaluation of reaching accuracy and variability, terminal feedback was provided every 6 trials. **C**. Action monitoring task. Participants next performed the goal-directed reaching movement that could either be non-deviated (control trials), or deviated by up to 30°. After ten non-deviated reaching and 12 practice trials, participants performed a block of 88 trials with randomized deviated and control trials. Based on this MA thresholds were determined. Once both tasks had been completed with one effector, they would be repeated with the other effector.

**Figure 2.**
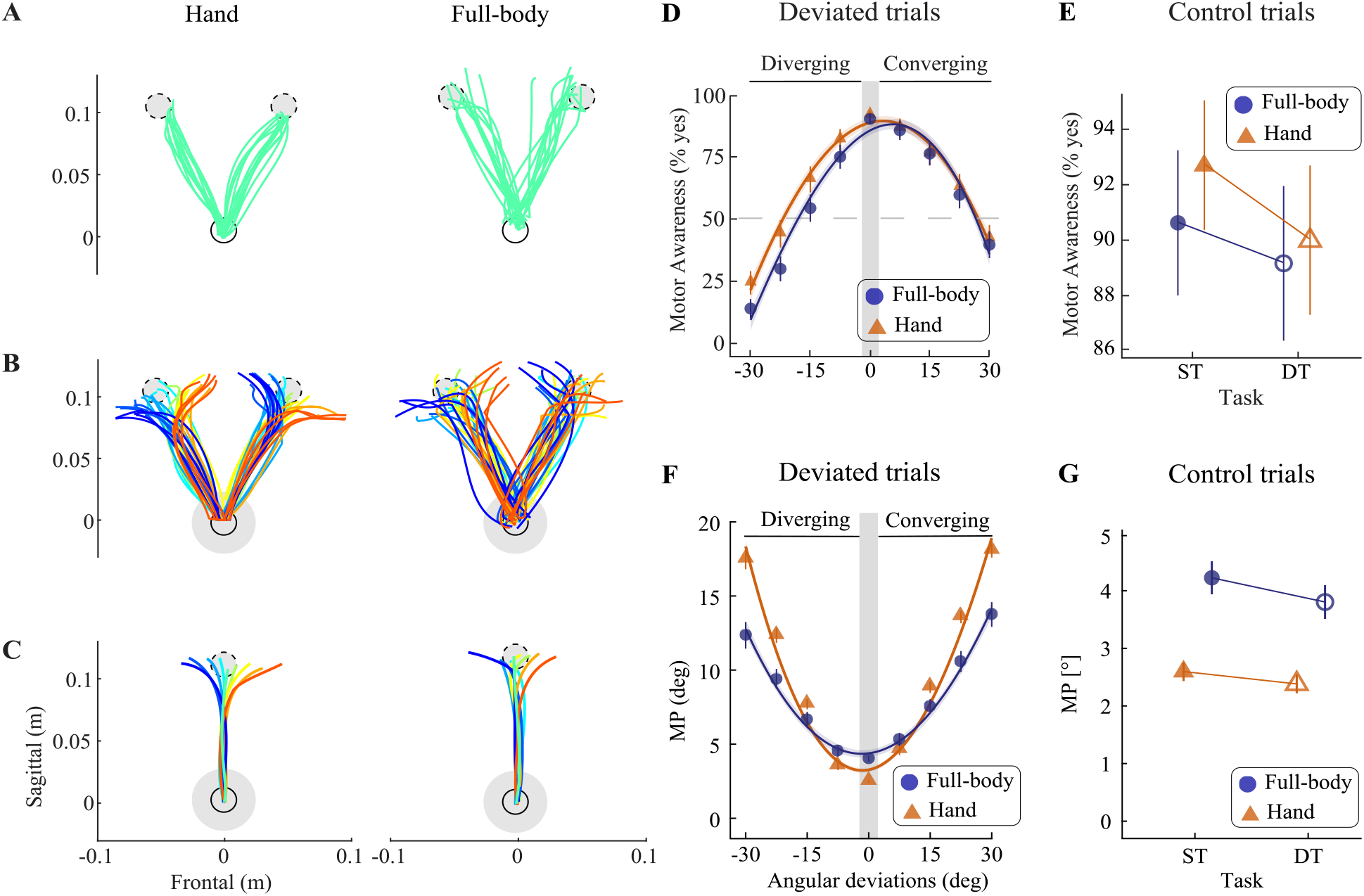
Motor Performance. A-C. Reaching trajectories of a single representative participant during the visually guided MA task. **A**. Trajectories from the resting position (solid circle) towards the two target locations (dashed circle) in control trials were more accurate for the hand (cf. panel G). **B**. Trajectories towards the targets in trials with deviations to the left (red tone lines) and to the right (blue tone lines). The shaded blue circle indicates the distance at which the deviations start. **C**. Average trajectories of this participant collapsed across the two target locations. **D**. MA was modulated by deviation-angle as well as the interaction of target- and deviation-angle. Participants made more attribution errors when the deviation angle converged with the target angle. **E**. Participants reliably judged feedback in control trials to match their own movement. **F**. Averaged MP across participants for full-body (round markers) and hand (triangle markers) for each deviation. **G**. Averaged MP across participants for non-deviated trials in single and dual tasks.

**Figure 3.**
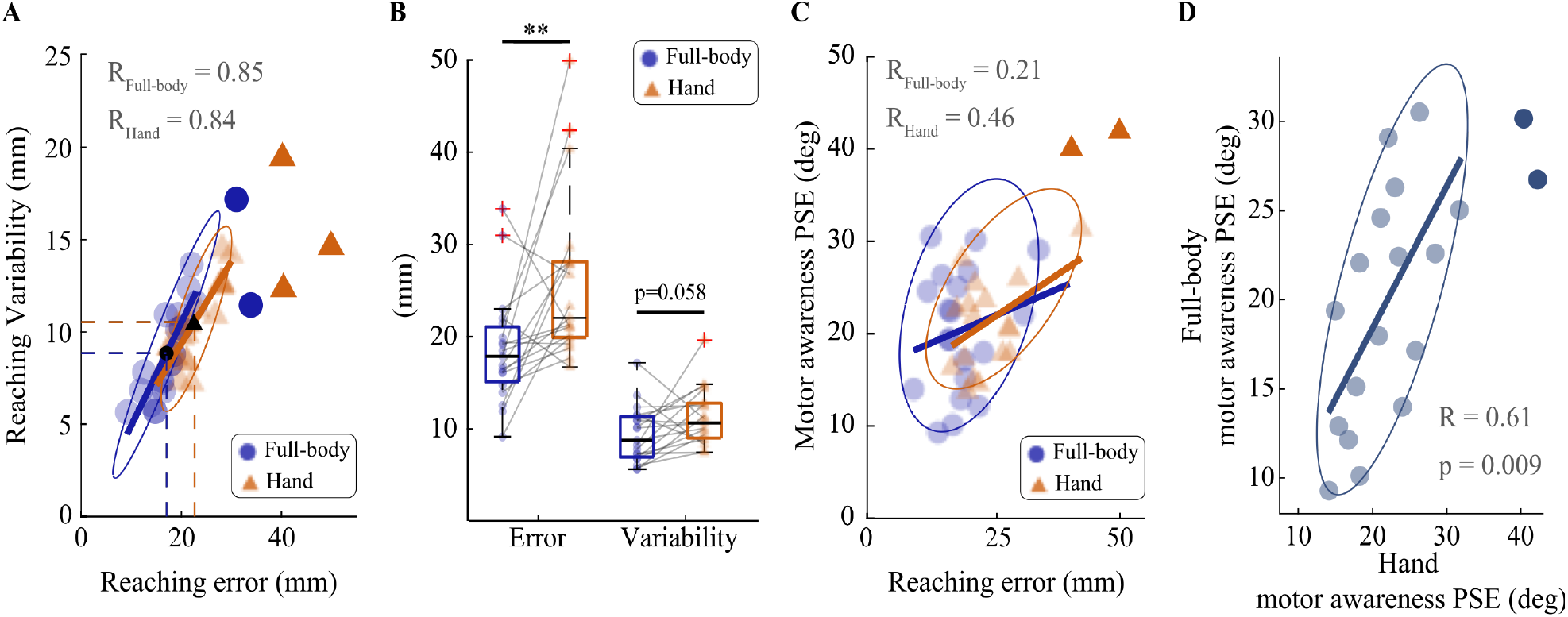
A. Blind reaching error and variability of each participant and for the full-body (blue dots) and the hand (orange triangles). Samples were obtained by averaging the 60 blind reaching trials of each participant. Opaque samples represent outliers, ellipses are the robust data centers and shaded areas are the 95% bootstrapped confidence interval. B. In average (back markers), blind reaching with the full body leads to more accuracy (smallest reaching error) and more precision (lowest variability), compared to blind reaching with the hand. C. Association between reaching error and motor awareness. Plotting symbols and shading conventions are the same as in A. D. Psychometric thresholds (point of subjective equality) for full-body and hand motor awareness show a strong positive correlation within participants.

All participants clearly distinguished between the introduced deviations, as demonstrated by the significant effect of Deviation (p<0.001, see Figure 2D). With respect to the Mode of the feedback, MA was significantly lower, that is more sensitive to deviations, when these were diverging (p<0.024). MA was also more sensitive for the full body compared to the hand (Effector, p<0.001) and during the dual task in general (p=0.014). Finally, a significant interaction between Mode and Effector (p=0.001) was detected. This interaction was driven by a significant difference in diverging trials between the two effectors, MA was more sensitive for full body movements.

We further extracted the psychometrical threshold, that is the point of subjective equality (PSE) for the detection of the introduced deviation. We first evaluated the main effect of Mode for converging versus diverging trials. A significant effect (p<0.001) was observed on the PSE, which was lower for diverging trials (cf. Supplementary Figure 1 A, inset). While the PSEs were similar for converging trials (28.1±2.34° for the hand and 26.9±2.33° for the full-body, mean±sem, BF_10_ =0.25, paired T-test: p = 0.63), they were significantly better for full-body movements (16±1.47°, against 19.9±2.04° for the hand, mean±sem, T-test: p=0.018) during diverging trials. Full-body and hand PSEs showed a strong positive correlation across participants (Figure 3D) as participants with higher thresholds for the hand also displayed higher thresholds for the full-body (Pearson r=0.6, p= 0.002).

Finally, we investigated whether there was an association between the participants’ ability to monitor their action, as expressed by the PSE, and their reaching accuracy derived in study 1 (Figure 3.C). This could account for potential differences observed in MA. However, Pearson’s skipped correlation was r=.21 (p=.4) for the full-body and r=0.46 (p=.46) for the hand, which represent low association factors.

### 2.2 Motor performance

Figure 2A shows the trajectories of the hand and full body in the sagittal plane from the resting position to each of the two targets for non-deviated trials. Visually guided reaching performance for non-deviated trials, independent of the task, was similar across participants and significantly better for the hand, contrary to the blind reaching data (Main effect of Effector, p<0.001, error_hand_=2.5°±0.06; error_full-body_=4.0°±0.1, mean±sem, see Fig. 2F grey shaded area). Somewhat surprisingly, we observed a significant reduction in reaching error during dual tasking independent of the effector (main effect of Task, p=0.01). There was no interaction between factors Task x Effector for non-deviated trials (p=0.38), see Fig. 2G.

All participants adapted their reaching movements to account for the target and angular deviations (Fig. 2B). Thus, trajectories were deviated in the opposite direction of the angular deviation, as reported for goal-directed tasks for the hand ^3,46,47^ and while walking ^5,31^. This compensation significantly increased with the experimentally induced angular deviation (main effect of Deviation (p < 0.001, Fig. 3F). MP was also significantly higher when the compensation was towards the center, i.e., when the deviation side converged towards the target side (see Supplementary Figure 1B). Therefore, a main effect of Mode was detected (p<0.001), as well as main effect of Task (p<0.001). As for control trials, dual tasking significantly improved motor performance. An interaction between Deviation and Effector was noticed (p<0.001). Motor compensation was higher for the full body during the smallest deviation (which may reflect lower accuracy during visually guided movements), while motor compensation was better for the hand during larger deviations (>15°).

### 2.3 Blind Reaching

We evaluated blind reaching performance as a baseline performance metric between effectors. The blind reaching error, representing reaching accuracy, was on average 26±9 mm (mean±std) for the hand and 18 ± 6 mm for the full body, while the effectors presented a variability, i.e. precision, of 11 ± 3 mm and 9±3 mm respectively (Figure 3A). These two metrics are highly correlated for the two effectors (Pearson’s skipped R > 0.84) as participants with small reaching errors (accuracy) also had displayed low reaching variability (precision). The average blind reaching error was significantly lower for the full-body than for the hand (paired T-test, p=0.007). Although reaching variability was higher for the hand than for the full-body, this difference did not reach significance (paired T-test, p=0.058) (see Figure 3B). The average reaching time across trials was shorter for the full-body than for the hand (paired T-test, p=0.007).

## 3 Discussion

A central challenge towards understanding our sense of agency from a sensorimotor account has been to quantify to what extent we are, or can be, aware of the details of our movements ^34,48^. Evidence from seminal case reports ^e.g. 49^ as well as behavioral studies has pointed to a dissociation between our movements and our awareness thereof ^50,51^, even when prompted to consciously monitor these (cf. introduction). Given the similar MA thresholds observed across tasks, effectors, as well as feedback modalities, it has been argued that there is an underlying, supramodal, and effector-independent monitoring mechanism ^33^ underlying our sense of agency. To test this hypothesis, we here assessed MA thresholds for goal-directed hand- and trunk movements within a single group of participants. Based on our findings, we propose that the central nervous system does make use of a general strategy to monitor goal-directed movements of our hand or the trunk, and that this is in the majority of cases based on abstract movement representations, rather than a direct comparison of effector-specific sensorimotor predictions and their reafferents. In-line with this, our results support the effector-independent hypothesis as we observed corresponding changes in MA depending on kinematic task demands (Fig. 2D), under cognitive loading, as well as a strong co-variance of MA thresholds within participants (Fig. 3D). That said, we also observed effector-specific differences, notably in diverging trials, indicative of a shift in strategy in case of kinematically more demanding tasks. We discuss our findings in relation to effector-independent representations in the motor domain and how such abstracted representations may play a role in understanding MA as the key sensorimotor component of our global sense of agency and the experience of the self as an intentional agent.

In the current study, we made several observations that support an effector-independent mechanism underlying our sense of agency. To begin with, SoA ratings in control trials matched across both effectors and conditions, suggesting that there are no differences for SoA at baseline, which roughly translates to everyday situations without significant external perturbations. The fact that we observed differences in motor performance for blind reaching, where full-body movements were more accurate and precise, and visually guided reaching in the MA task, where hand-movements were superior, gives an indication of the inherent tolerance of the mechanism and links to the aforementioned dissociation between action and perception. These results also extend our findings for goal-directed walking ^5,44^ and continuous walking ^4^ where, as observed in the current tasks, MA ratings in control trials were not affected by cognitive loading, independent of its detrimental effects on motor performance ^41,42^.

Further evidence for effector-independence comes from deviated trials, where body-part and full-body MA thresholds showed a strong correlation within participants: Participants that were better at detecting experimental deviations of their hand-movements, also picked up on more of the same deviations during full-body movements. In other words, there is correspondence for the tolerance in the monitoring mechanism within participants. Extending this similarity, we observed clear and corresponding effects of kinematic task demands on MA ratings for the hand and full body in converging trials, where participants had to compensate for the experimental deviation by moving towards a central location, straight ahead. When the experimental deviation “aided” the movement in this way, MA thresholds significantly increased for both effectors. That is, fewer mismatches were detected. While these two findings are in-line with an effector-independent mechanism, the fact that they persist even though we observe effector-specific differences in motor compensation may point to a more general commonality, namely that of an abstract motor representation serving as a basis for the monitoring mechanism. As we observed, particularly for walking under cognitive loading ^5,44^, conscious monitoring, unlike the continuous automatic adjustment observed for sensorimotor control, may rely on a movement representation formed in the motor planning phase before the onset of the movement.

As previously proposed ^33,52^, a goal-directed paradigm in itself may create a type of bias linked to the successful completion of the task as determined at the planning stage: MA ratings are consistently higher for successfully completed trials even if sensorimotor re-afferents about the on-going movement contain a strong error signal ^32^. In addition to the “limits of conscious action monitoring” described in ^53^ and already evident in ^11^, the evidence for such a bias when comparing converging and diverging trials goes beyond simple visual dominance ^54–56^.

What happens when the combination of the target location and the introduced deviation make it more difficult to reach the target? Although we report matching MA thresholds in non-deviated and converging trials, independent of underlying sensorimotor differences, MA ratings differed in the case of diverging trials (Figure 3D and SFigure 1A). In this case, the introduced deviation is towards the inside of the target, i.e. the midline, such that compensatory movements must be made to the outside, i.e. the periphery. As it became kinematically more challenging to complete the task, MA ratings became increasingly more sensitive to the introduced deviations. MA thresholds were significantly lower, indicating that participants were more likely to reject even slightly deviated trials. This effect was evident across both effectors, but especially pronounced for full-body movements. Overall, this change in MA thresholds in diverging trials is not surprising. As the motor task becomes more difficult, reaching accuracy decreases, meaning that visual errors at the end point are larger, on top of the increasingly conflicting proprioceptive reafferents. As the target is not clearly reached, even with increased effort, there may be less incentive to consolidate this conflicting information.

The more pressing question is why differences in MA thresholds between full-body and hand movements appear in these trials. As before, the answer does not appear to primarily lie with participants’ motor performance. The difference observed in motor performance was also evident in converging trials. It appears then that in case of exacerbated sensorimotor conflict, the proposed effector-independent mechanisms may be augmented, or reweighted, with respect to effector-specific sensorimotor information similar to a reliance on prior knowledge under uncertainty ^57^. Two possible explanations come to mind. For one, in the case of full-body movements the vestibular system may provide an additional, more relied upon cue of heading information, in particular when the deviated visual feedback becomes less reliable ^58^. This is in-line with the improved reaching accuracy and lower variability we report for blind-reaching with the full-body ^see also 45^. As this non-visual representation of the full-body movement appears to be more accurate than its hand-equivalent, it may more strongly influence MA ratings in diverging trials leading to the differences between effectors. For another, and tying into this explanation, such a reweighting would also be indicative of a shift in monitoring from an abstract motor representation involved in motor planning, to a stronger reliance on effector-specific reafferent feedback during motor execution ^59^, which as recent evidence suggest may be implemented simultaneously ^60^. Future studies could investigate such a reweighting systematically by changing the reliability of visual, vestibular, and proprioceptive information when assessing MA ^58,61^.

The notion of abstract motor representations is not a new one. Theories of sensorimotor control have long considered effector-independent components across actions based on concepts such as “motor primitives” and “motor equivalence” ^8,9,62^. Therefore, paradigms that put an emphasis on the content of the movement itself provide a unique opportunity to study the hypothesis that MA and the sense of agency rely on similar abstract motor representations. In their paper on an effector-independent action system, Liu and colleagues ^7^ discuss the example of letters written with different body parts ^62,63^ as a further example of motor equivalence (see also the accompanying commentary by Goodale ^64^). This is similarly interesting from a conceptual standpoint for MA as e.g., writing the letter “a” with your foot, although now more closely tracked than for a simple reaching movement or while writing the letter with your hand, will still be evaluated to the notion of the letter “a” rather than an explicit sensorimotor transformation. Indeed, this was observed in experiment 3 of ^13^, where figural discrepancies significantly increased awareness of sensorimotor mismatches compared to velocity mismatches.

Evaluating continuous movements that are not inherently goal-directed may provide further insights into this characteristic of MA and the sensorimotor components underlying our sense of agency. In these studies, the movement itself is monitored, rather than a pre-specified outcome. The cyclical nature of movements such as opening and closing the hand ^65^, drawing ^13^, walking ^4,6^, or breathing ^66^ already demonstrate examples of automatic sensorimotor synchronization that can coincide with or oppose MA ratings. Teasing out such quantifiable differences, how they may be abstracted in terms of motor primitives, equivalence, physiological requirements, figural descriptions, or explicitly desired states will be required to develop a clearer picture of the level of conscious access to our motor system as well as the sensorimotor contributions to our SoA.

Although the link between “low-level” sensorimotor control and “high-level” cognitive intentions and their importance for our sense of agency is intuitive and has long been the subject of research, conceptual accounts ^48,67^ and computational models, originally designed purely for sensorimotor control ^68,69^, have thus far failed to provide a coherent framework. Notably, this issue arises at the level of abstraction, such that explanations have not been extrapolated from “inaccessible” sensorimotor control ^48^ or relied on descriptive “top-down” and “bottom-up” interactions between categorically different types of representation ^67,see also 70’s hierarchical model for error detection^. Our current findings provide a common denominator for these approaches by showing that MA may well depend on sensorimotor transformation, as in the case of conflicting feedback, but additionally makes use of effector-independent movement abstraction already observed in the motor system. This “Ignorance of the effector systems” ^71^ has also been proposed as a prerequisite to understanding authorship and ownership for our thoughts in terms of internal models or predictive coding ^37,72,73^.

### 3.1 Conclusion

What do we have in mind when we identify an on-going movement as our own, when we feel to be authors of our own actions? While previous frameworks have alternated between categorically different, body-part-specific sensorimotor transformations on the one hand to abstract intentions on the other, we propose that motor awareness and our Sense of Agency for our movements generally rely on effector-independent representations or motor-abstracts. If there is a mediated action, as for example when flipping a light switch, its outcome may override internal representations about the actual movement. Similarly, under situations of high-uncertainty or difficulty, we may have to focus on effector-specific transformations. However, most of our movements do not fall into these categories, yet we maintain a reliable sense of control over them and perceive correspondence (or at least no conflict) with our movement plans. Understanding MA as an effector-independent process that draws on abstract motor representations provides a first step, a proof by induction, to linking primary sensorimotor transformations and motor actions to abstract representations of our intentions and thoughts.

## 5 Materials and methods

### 5.1 Participants

Twenty-two healthy adults were enrolled for this study (15 females, mean age = 25 ± 3.9 years, height = 1.66 ± 0.07 m). The sample size was determined based on the large effect size of the spatio-temporal mismatches reported in prior studies ^4,6,23,74^. This sample is large enough to be indicative of a dual-task effect and motivate an appropriately confirmatory study if desired. Due to the wide range of secondary tasks available and mixed findings across different participant cohorts, it is difficult to determine a meaningful sample size a priori.

Participants had intact or corrected to normal vision, intact musculoskeletal system, and no history of orthopedic, neural or psychiatric conditions. The experimental procedure described in this study was approved by the local ethics committee in accordance with the ethical standards laid down in the 1964 Declaration of Helsinki.

Two participants were discarded from the data analysis. The first one could not complete the balance task since they felt dizzy. They were the only volunteer to report vertigo. The second exclusion was done post-analysis, since the motor awareness threshold of this participant was below the point of subjective equality for all deviations, including control trials with no deviation.

### 5.2 Experimental setup

For the full-body configuration, participants were secured via straps at the pelvis level inside the THERA-Trainer coro ^75^, a therapy device for dynamic balance exercises used to track the movement of the full-body. Setup takes approximately 3 minutes and participants are securely positioned in a standing position, preventing potential falls. Its frame is height-adjustable and composed with two spring elements, whose resistance was set to minimal, for this study. The behavior of the spring system is comparable to the one of a joystick, meaning that it always pulls back towards the center. No degrees of freedom were locked during the experiment. It is equipped with an inertial measurement unit (IMU) which wirelessly transmits the tilt of the frame at a sampling rate of 30 Hz. The projection of the center of mass (CoM) in the frontal and sagittal direction was estimated by multiplying the pelvis height with the sinus of the roll and pitch angles given by the IMU, respectively. The upper frame of the THERA-Trainer core was modified to host a Logitech ATTACK 3 joystick, whose position was also streamed at 30 Hz.

### 5.3 Protocol

This study was divided in two parts. It started with 1) a blind reaching experiment, where participants were asked to reach a target without any visual feedback, and 2) an action monitoring study allowing the assessment of participants’ MA threshold. The order of the studies were not counterbalanced to not bias the near-space representation with the visuomotor conflicts present in the action monitoring task.

#### 5.3.1 Study 1: blind reaching

This study was composed of two experimental blocks; a full-body and a hand blind reaching block effectuated in a randomized order. A cursor representing either the projection of participant’s CoM or the hand position in the transversal plane was displayed on LCD monitor. Each trial started from the center target representing the resting position (i.e. standing straight or center of the joystick). A virtual target appeared on the top of the screen at one of two randomized locations. The two target locations were placed at ±25° from medial axis along a circle with a radius of 80% participant’s range of motion. Participants were asked to perform a hand movement with a joystick or to lean with their full-body to reach the target. During the full-body block, they were instructed to keep the feet shoulder-width apart, to maintain a static base of support. For the hand block, they used their dominant hand.

All trials begin from the resting position. The experiment started with a training phase where they were asked to reach one of the two virtual targets with congruent visual feedback. In the second phase (blind reaching learning), the cursor disappeared once the virtual target was presented. Participants were asked to reach blindly the virtual target, stop and stay once they think they reached the virtual target. They received terminal visual feedback of the cursor position to show them their reaching accuracy. They had 20 trials to improve their reaching accuracy. In the last phase, the terminal visual feedback appeared every 6-trial only.

#### 5.3.2. Study 2: action monitoring

Participants performed two experimental blocks, a full-body (FB) and a hand (H) action monitoring block, effectuated in a randomized order. These two blocks were divided in two sub-blocks, comprising a single task (ST) and a dual task (DT). The sub-blocks order was randomized within blocks. The experimental procedure is illustrated in Fig. 4B.

**Figure 4:**
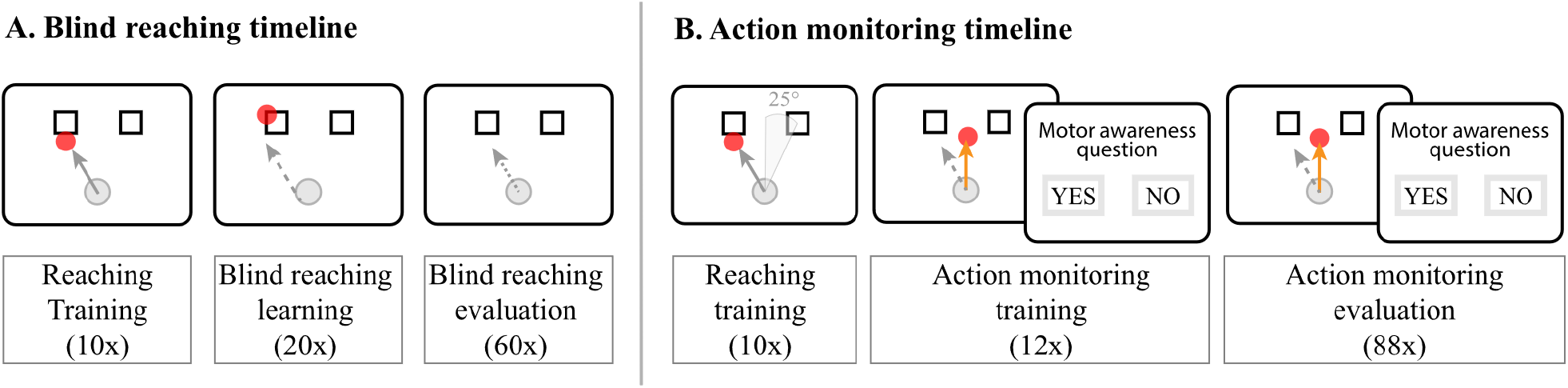
Timelines of the tasks. Panel A.A. Timeline of the blind reaching task. The effector order was randomized across participants. Participants started the experiment with a reaching training with a concurrent and congruent visual feedback to familiarize themselves with the setup. They were then trained to reach the target with only a terminal feedback. During the evaluation, they had a terminal feedback every 6 trials. Panel B. Timeline of the action monitoring task. Participants started in a randomized fashion either by controlling the cursor with the full-body or the hand. The order of the tasks (single vs dual task) with which they started was also randomized. The experiment started with a familiarization phase, then participants learnt to reply to the motor awareness question, and finally their MA threshold was evaluated. Once they completed both tasks with one effector, they repeated the experiment with the other effector.

Our action monitoring task was similar to the paradigm used in previous studies to test for motor awareness through perceptual visuomotor conflicts of voluntary movements ^53^. As in study 1, they were able to control a cursor either with their hand or by tilting the body. In both cases, they were asked to produce a trajectory as straight and smooth as possible. After reaching the target, they returned to the resting position to start the next trials. Return movements to the resting position were not analyzed, since they are assisted by the spring system of the THERA-Trainer coro or of the joystick.

The experiment started with a training phase (20 trials) where participants familiarized themselves with the devices and the task. The cursor position was congruent with their movement. Participants then performed the action monitoring training (12 trials) and evaluation (88 trials). In some trials (75%) and beyond a distance of 250 mm from the resting position, the trajectory of the cursor was deviated either clockwise (+ sign) or counterclockwise (-sign) by an angular rotation of 7.5°, 15°, 22.5° or 30°. The amplitude and the side of the deviation was randomized and evenly distributed across targets (88 trials per sub-block, including 24 control trials, i.e. no deviations, and 16 trials per deviation) as in ^31^. At the end of each trial, participants used the joystick for both blocks to answer a question displayed on the screen asking: “Did the movement you saw on the screen correspond to the movement of your hand/body?” ^23^.

In the dual task sub-blocks, participants performed the same action monitoring task, while executing a visual color-word stroop task ^76^. A color name (e.g. BLUE printed in red) appeared inside the cursor along the trajectory at 8 different distances from the resting position, so that participants could not predict the onset. The font color always conflicted with the color name. Participants were asked to say the font color out loud as fast as possible and then reply to the MA question quoted above.

### 5.4 Data analysis

Data were processed offline using Matlab (MathWorks, Natick, Massachusetts, USA) and the R environment ^77^. For both tasks, we first analyzed the reaching trajectories. Those were smoothed with a 5-window moving average filter, interpolated and finally averaged across targets and deviations for each participant. Trials where participants restarted their movement (i.e. came back to the center and reached again) were excluded.

#### 5.4.1 Study 1: Blind reaching

The reaching error of each trial was defined by calculating distance between the trajectory end point and the target center. The error amplitude is a direct measure of the spatial accuracy. To quantify the overall accuracy of each effector, reaching errors were then averaged across participants. To evaluate the participants consistency in their performance, the reaching variability was calculated by taking the standard deviation of the reaching errors and averaged across participants to assess the precision of each effector.

#### 5.4.2 Study 2: action monitoring

First, we analyzed the responses to the action monitoring question. Motor awareness (MA) was defined as the number of yes-responses out of all trials of the same deviation. Correct self-attribution or MA was a “yes” response for non-deviated trials and a “no” response for deviated ones. The MA threshold was determined by fitting a psychometric curve to the participants’ responses across the absolute delays and extracting at the 50% point of subjective equality (PSE). PSE was then averaged across participants.

Second, we measured the trajectory endpoint to assess motor performance (MP). The trajectory endpoint corresponds to the effector position when reaching the target radial distance, which is not equivalent to the trial endpoint (i.e. center of the target). MP was defined as total angle compensated by the participants considering the endpoint of each movement trajectories and measured from the onset of the deviation, which was 25 mm from the resting position.

Finally, as additional variables, we analyzed the reaching and response time. Reaching time was calculated from movement onset to end of the trial. Movement onset was defined by the radial velocity being greater than 2% of the maximum trajectory velocity. The response time corresponds to the time from the appearance of the question to the participant’s response via joystick button. Although we recorded response times, participants were not asked to reply to the motor awareness question as quickly as possible. They also could move to target at a speed that was comfortable for them.

The aforementioned dependent variables were analyzed as functions of the independent variables such as Deviation, Deviation Side, Target Side, Task, Mode, and Effector. Deviation Side was defined as the direction of the deviations (i.e. clockwise or counterclockwise). The target side was the position of the target with respect to the midline (left or right). The mode defines if the deviation converged towards or diverged away from the target position.

### 5.5 Statistical analysis

Mixed-effects linear models were conducted on MA (logistic), MP, and the two observatory variables as well as the PSE (see supplementary table 1). For MA, the first model evaluated the effect of Target Side on control trials (m1). Then, we tested the interaction Task x Effector also on control trials (m2). On deviated trials, first the interaction Target Side x Deviation Side was tested (m3). Then, deviations were split in two modes (converging and diverging). We tested the effect of Deviation (m4) and then their absolute values were taken. We tested the interaction Deviation x Mode as well as the effect of Effector and Task (m5). Finally, for our last model, we tested the interaction Mode x Effector (m6) since this was not possible in the previous model. Models m1 to m4 were applied to MP to evaluate the effects of independent variables. The interaction Deviation x Effector and the effect of Mode and Task were evaluated in another model (m5’) For all the above-mentioned models, participants were specified as a random (subject) factor, allowing for random slopes and intercepts.

Paired Student’s and Bayesian t-tests were used to determine whether the distributions of yes-responses for control trials were significantly the same for Effector and Task, as well for the MA PSE. Bonferroni’s correction was applied for multiple comparison. Paired Student t-tests were used to assess significant differences between full-body/hand reaching accuracy and variability. For correlations analysis, we use a robust procedure called Pearson’s skipped-correlations ^78^. This method protects against bivariate outliers. We measured the strength of the association between average reaching error and PSE of each participant.

## Supplemental Information

### Supplemental Results

#### Response Times

We observed a significant difference in response times depending on the Effector (p<0.001) as participants answered significantly faster when the effector was the hand. Response times were not significantly affected by the magnitude of the deviation (p=0.34).

#### Reaching Completion Times

Reaching times for control trials were significantly shorter for the hand than for the full body (Main effect of Effector, p<0.001). Dual tasking led to shorter reaching times for the full-Body, whereas they remained similar for hand movements (interaction Task x Effector, p=0.001).

Reaching time significantly increased with the introduced deviation (Main effect of Deviation, p<0.001). As for control trials, reaching times were significantly higher for the full body than for the hand (main effect of Effector, p<0.001). Dual tasking significantly decreased reaching duration (main effect of Task). The reaching time was lower when the deviations were converging (main effect of mode, p < 0.001).

#### Weak Relationship between Blind Reaching and MP

The association between the participants’ MP and their reaching accuracy derived in study 1 was assessed. Pearson’s skipped correlations were lower than 0.32 for the full-body and lower than 0.04 for the hand, which represent low association factors.

#### Dual Task Performance

Finally, during the dual-task, participants named the wrong color of the target on average 0.74±0.2 times (mean ± sem) over 88 trials. No significant difference was detected for the two effectors (paired T-test, p=0.71).

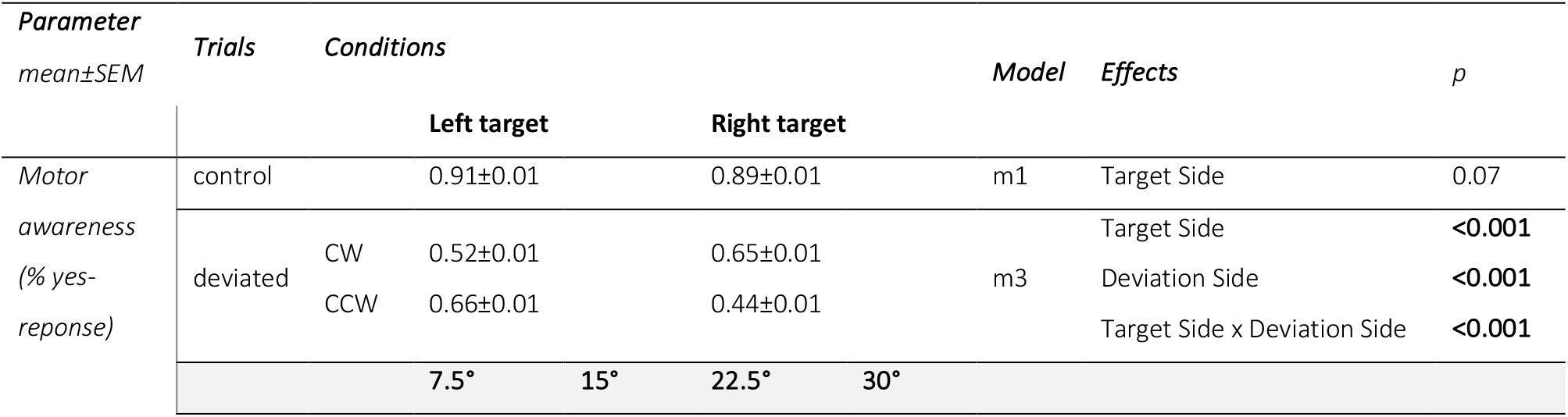
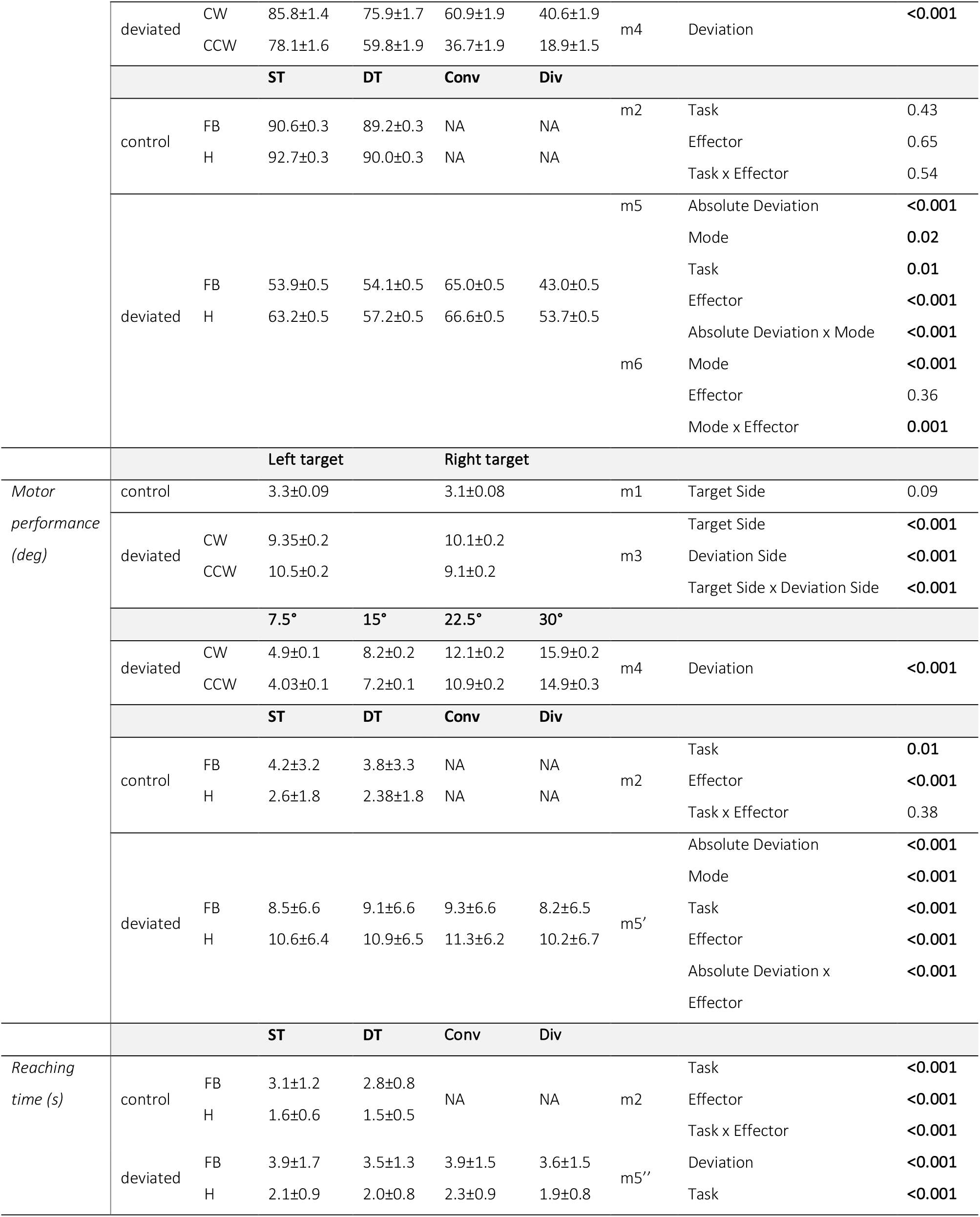
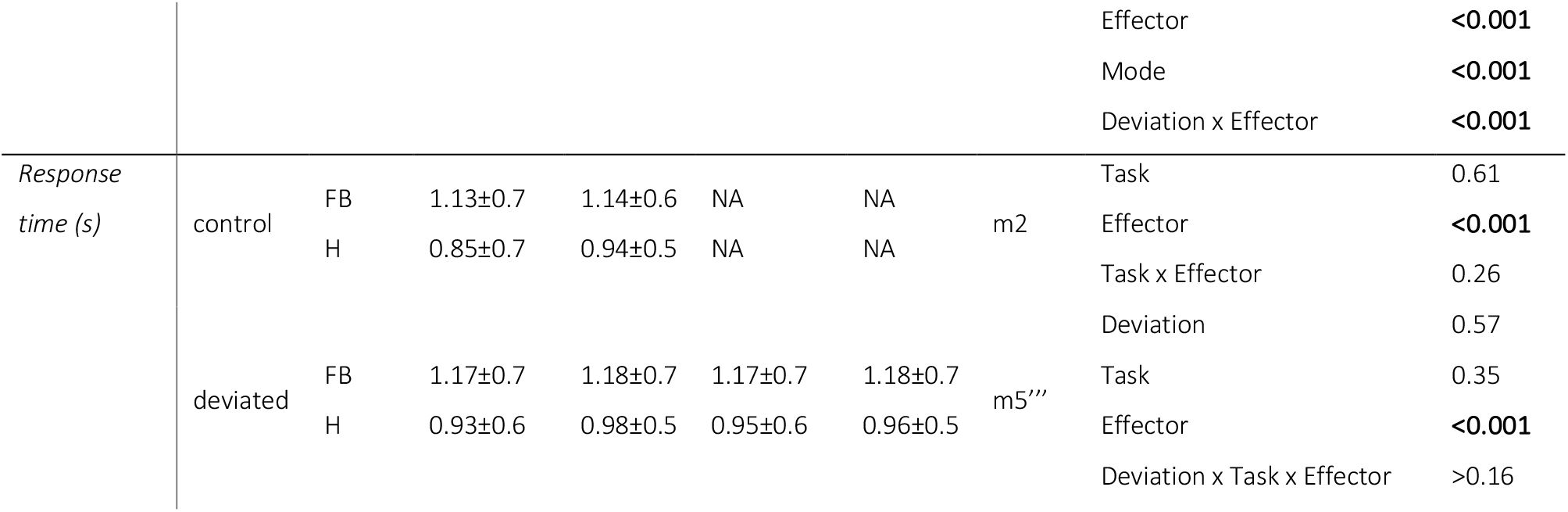

**SFigure 1:**
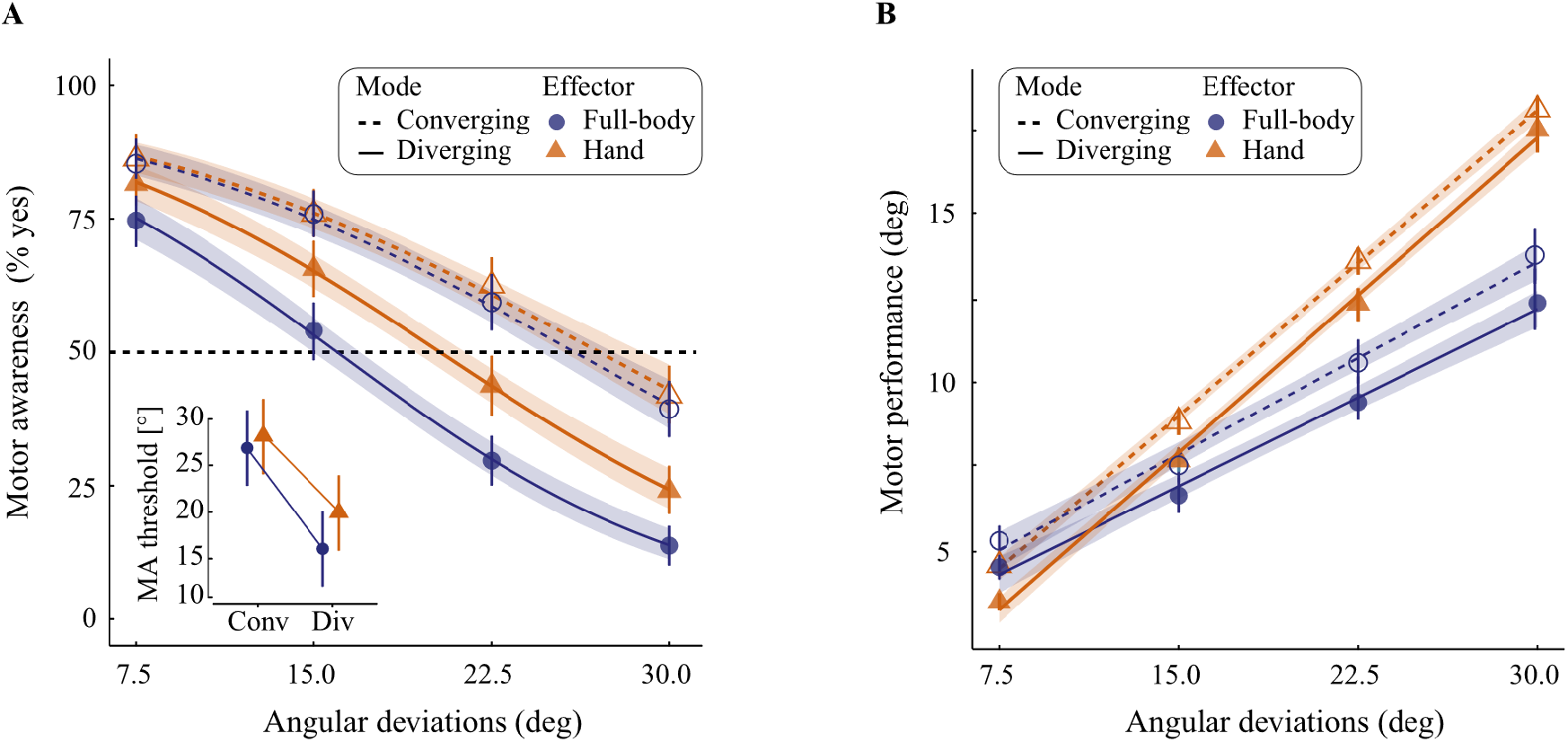
Effect of kinematic demands on motor awareness (A) and motor performance (B). The two effectors have similar motor awareness thresholds for converging trials, while MA threshold was better for full-body during diverging trials. Motor performance is better for the hand and during converging trials.

